# NOS2 and COX2 Blockade Limits TNBC Disease Progression and Alters CD8^+^ T Cell Spatial Orientation and Density

**DOI:** 10.1101/2022.06.03.494733

**Authors:** Veena Somasundaram, Lisa A Ridnour, Robert YS Cheng, Abigail J Walke, Noemi Kedei, Dibyangana D Bhattacharyya, Adelaide L Wink, Elijah F Edmondson, Donna Butcher, Andrew Warner, Tiffany H Dorsey, David A Scheiblin, William Heinz, Richard J. Bryant, Robert Kinders, Stanley Lipkowitz, Stephen TC Wong, Milind Pore, Stephen M. Hewitt, Daniel W McVicar, Stephen K Anderson, Jenny Chang, Sharon A Glynn, Stefan Ambs, Stephen J. Lockett, David A Wink

## Abstract

Anti-tumor immune polarization is a key predictor of clinical outcomes to cancer therapy. An emerging concept influencing clinical outcome involves the spatial location of CD8^+^ T cells, within the tumor. Our earlier work demonstrated immunosuppressive effects of NOS2/ COX2 tumor expression. Here, we show that NOS2/COX2 levels influence the polarization and spatial location of lymphoid cells including CD8^+^ T cells. Importantly, elevated tumor NOS2/COX2 correlated with exclusion of CD8^+^ T cells from the tumor epithelium. In contrast, tumors expressing low NOS2/COX2 had increased CD8^+^ T cell penetration into the tumor epithelium. Consistent with a causative relationship between these observations, pharmacological inhibition of COX2 with indomethacin dramatically reduced tumor growth of the 4T1 model of TNBC in both *WT* and *Nos2*^*-/-*^ mice. This regimen led to complete tumor regression in ∼20% of tumor-bearing *Nos2*^*-/-*^ mice, and these animals were resistant to tumor rechallenge. Th1 cytokines were elevated in the blood of treated mice and intratumoral CD4^+^ and CD8^+^ T cells were higher in mice that received indomethacin when compared to control untreated mice. Multiplex immunofluorescence imaging confirmed our phenotyping results and demonstrated that targeted Nos2/Cox2 blockade improved CD8^+^ T cell penetration into the 4T1 tumor core. These findings are consistent with our observations in low NOS2/COX2 expressing breast tumors` proving that COX2 activity is responsible for limiting the spatial distribution of effector T cells in TNBC. Together these results suggest that clinically available NSAID’s may provide a cost-effective, novel immunotherapeutic approach for treatment of aggressive tumors including triple negative breast cancer.

## Introduction

Breast cancer is the most common tumor globally among women and the leading cause of cancer-related deaths among women in North America (1, 2). It is a heterogeneous disease that can be classified as estrogen receptor positive (ER+) or the more aggressive ER negative (ER-) subtype. The upregulation of proinflammatory signaling pathways has been reported in breast cancer, suggesting a regulatory impact on the immune system (3). In ER- and triple negative breast cancer (TNBC), nitric oxide synthase-2 (NOS2) and cyclooxygenase-2 (COX2) have been described as independent predictors of disease outcome (4-6). Recently, elevated tumor coexpression of NOS2/COX2 demonstrated strong predictive power associated with poor survival among ER-patients as defined by a hazard ratio (HR) of 21 (7). In addition, NOS2-derived NO and COX2-derived prostaglandin E2 (PGE2) were shown to promote feed-forward NOS2/COX2 crosstalk where NO induced COX2 and PGE2 induced NOS2, as well as other downstream signaling targets (7). While NOS2-derived NO is a driver of oncogenesis, chemoresistance, metastasis, and immunosuppression, COX2 supports both NOS2 expression and mediates immune suppression both systemically and within the tumor microenvironment (TME) (7-11). Studies have shown that NOS inhibition improved treatment efficacy by down regulating IL-10 within the TME (12, 13). Importantly, a recent phase 1/2 clinical trial has demonstrated improved clinical outcome defined by an overall response rate of 45.8% in patients with locally advanced breast cancer (LABC) and metaplastic TNBC that received the pan-NOS inhibitor L-NMMA and aspirin combined with taxane (14). Importantly, this study also revealed that 27.3% of treated LABC patients achieved pathological complete response at surgery where remodeling of the tumor immune environment was observed in patients responding to this therapy (14). Given that COX2 promotes immune suppression within the TME, targeting both NOS2/COX2 may further improve outcome and thus provide a novel option in breast cancer treatment by augmented anti-tumor immune response (9, 10).

Recent studies have shown that the localization of CD8^+^ T cells provides important insight into tumor response and patient survival (15). Tumors with enhanced CD8^+^ T cell density and penetration into the tumor epithelium have improved outcomes compared to tumors where CD8^+^ T cells are absent, sparse, or spatially restricted to the tumor margin or stroma (15). Tumor expression of NOS2 and COX2 are predictive of ineffective immune response and poor outcomes, however how expression impacts the spatial distribution of immune cells remains unclear (4, 5, 7, 8).

Multiplex spatial imaging provides a powerful tool to improve our understanding of how the spatial localization of tumor cellular neighborhoods including the tumor immune microenvironment impacts clinical outcome. Here we spatially examined TNBC tumors and found that those high in both NOS2 and COX2 showed an immunosuppressed phenotype with reduced infiltrating CD8^+^ T cells that were restricted to the tumor margin or stroma. Modeling of TNBC in mice showed that the clinically available NSAID indomethacin substantially reduced tumor growth, restored T cell numbers and resulted in their improved spatial distribution within the tumor. This work proves that COX2 activity is responsible for limiting the spatial distribution of effector T cells in TNBC and suggests that NSAID’s may provide a cost-effective, novel immunotherapeutic approach for treatment of aggressive tumors including TNBC.

## Results

### NOS2^*Lo*^/COX2^*Lo*^ Tumors exhibit enhanced T cell infiltration

Tumor NOS2/COX2 co-expression predicts poor clinical outcome via promotion of breast cancer disease progression by several mechanisms including altered immune signaling (7-12, 14). Herein, we explored the impact of abated NOS2/COX2 signaling on 1) immune polarization, 2) the spatial location of immune mediators, and 3) tumor response and outcome. Immune status was evaluated using multiplex imaging in 26 ER-breast tumors previously examined for NOS2/COX2 expression (4, 5, 7). From these tumors, 16 were selected for spatial analyses based upon TNBC phenotype and NOS2^*Lo*^/COX2^*Lo*^ vs NOS2^*Hi*^/COX2^*Hi*^ tumor expression. Multiplex imaging was performed to evaluate CD8^+^ T cell infiltration and spatial location associated with NOS2^*Lo*^/COX2^*Lo*^ vs NOS2^*Hi*^/COX2^*Hi*^ tumor expression. Fluorescent antibodies targeting CD3, CD4, CD8, CD68, FOXP3, PDL1, PD1, together with KRT1 (CK-1) and SOX10 (CKSOX10) as tumor markers, were used to spatially define the immune microenvironment in 16 TNBC tumors. Fig. 1A demonstrates a clear infiltration of CD8^+^ T cells (red) into the tumor core (CKSOX10, blue) of NOS2^*Lo*^/COX2^*Lo*^ expressing tumors. In contrast, NOS2^*Hi*^/COX2^*Hi*^ expressing tumors exhibited markedly reduced CD8^+^ T cell infiltration, with stromal restriction as shown in Fig. 1B or marginal restriction as shown in Fig. 1C. In addition to stroma- and margin-restricted T cells, NOS2^*Hi*^/COX2^*Hi*^ tumors exhibited areas lacking immune cells (immune deserts, Fig. 1B white arrow), which has been associated with poor outcome (15). Fig. 1B also shows dispersed clusters of tumor cells (Fig. 1B white arrow), which may be indicative of more invasive tumor phenotypes that support poor clinical outcome as previously described (7). Importantly, cell quantification revealed that NOS2^*Hi*^/COX2^*Hi*^ tumors had significantly fewer infiltrating CD8^+^ T cells when compared to NOS2^*Lo*^/COX2^*Lo*^ tumors (Fig. 1D).

**Fig. 1.**
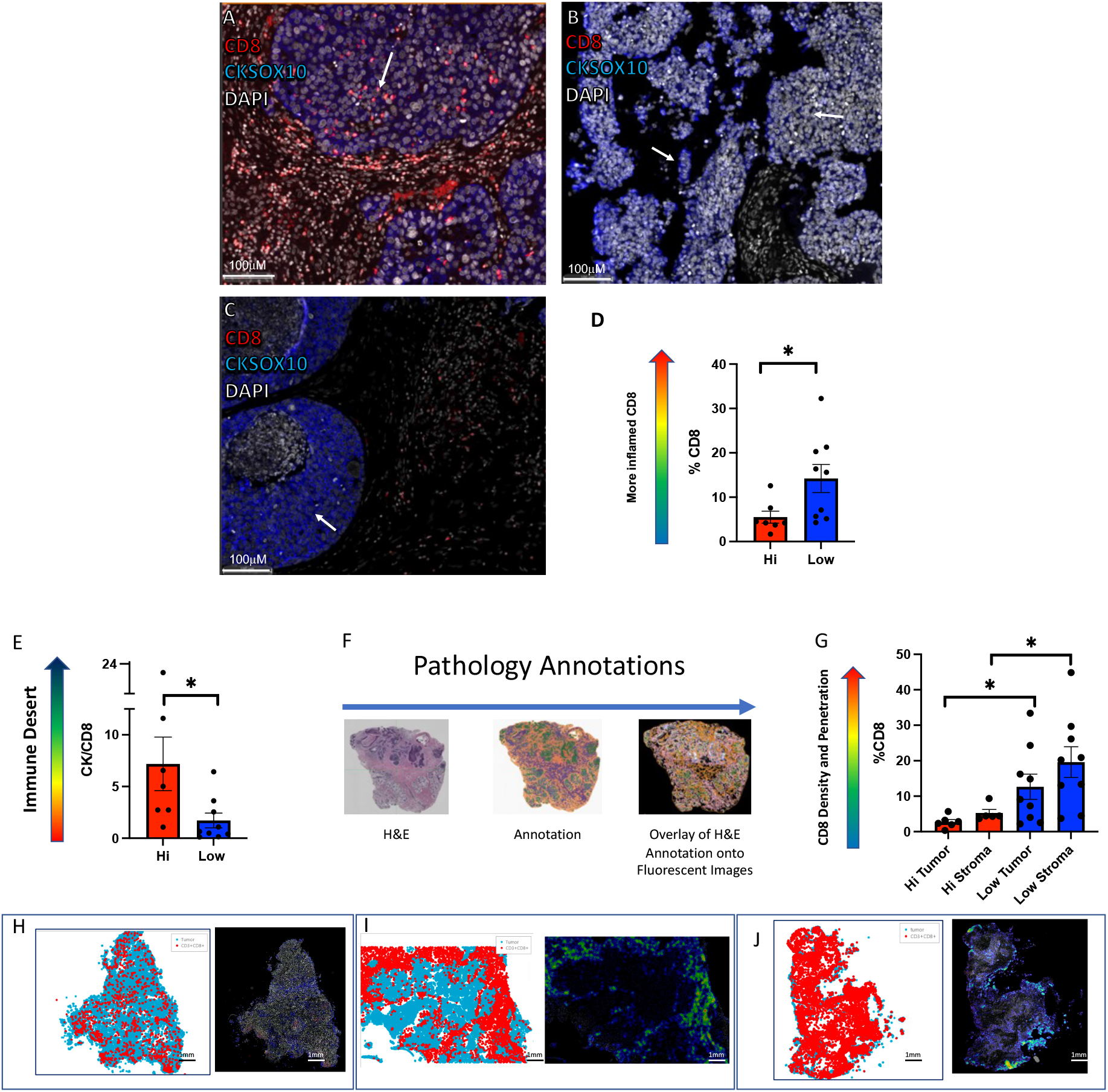
Multiplex Fluorescence Imaging evaluating altered CD8^+^ T cell infiltration in TNBC NOS2^Lo^/COX2^Lo^ vs NOS2^Hi^/COX2^Hi^ expressing tumors. Spatial distribution at 100μm magnification showing CD8^+^ T cells (red) relative to the CKSOX10 (blue) tumor marker in A) fully inflamed NOS2^Lo^/COX2^Lo^ tumor B) immune desert region in NOS2^Hi^/COX2^Hi^ tumor C) stroma restricted CD8^+^ T cells in NOS2^Hi^/COX2^Hi^ tumor. D) Percent CD8^+^ T cells are elevated in NOS2^Lo^/COX2^Lo^ vs NOS2^Hi^/COX2^Hi^ expressing tumors. E) Elevated CKSOX10/CD8^+^ T cell ratio in NOS2^Hi^/COX2^Hi^ vs NOS2^Lo^/COX2^Lo^ tumors. F) Pathology annotations for spatial distribution of CD8^+^ T cells in NOS2^Lo^/COX2^Lo^ vs NOS2^Hi^/COX2^Hi^ tumors. G) Quantification of %CD8^+^ T cells distributed in tumor vs stroma of NOS2^Lo^/COX2^Lo^ vs NOS2^Hi^/COX2^Hi^ expressing tumors. CD8 spatial distribution and density heat maps in whole tumor images at 1mm magnification showing H) fewer CD8^+^ T cells that are I) restricted to stroma as well as immune deserts in NOS2^Hi^/COX2^Hi^ tumors and J) densely infiltrated CD8^+^ T cells in fully Inflamed NOS2^Lo^/COX2^Lo^ tumor. (Significance determined by Mann Whitney two-tail test).

To further examine the influence of NOS2/COX2 tumor expression on CD8^+^ T cell infiltration, the ratio of the tumor markers CKSOX10 to the CD8^+^ T cell marker, which is a more direct regional tumor-to-CD8 comparison, was determined. The CK/CD8^+^ T cell ratio provides a metric of both CD8^+^ T cell infiltration and density where higher numbers represent tumors with reduced CD8^+^ T cell numbers associated with immune deserts. As seen in Fig. 1E, NOS2^*Hi*^/COX2^*Hi*^ tumors had significantly higher tumor CK/CD8^+^ T cell ratios when compared to NOS2^*Lo*^/COX2^*Lo*^ tumors. These observations support a role of elevated tumor NOS2/COX2 in the regulation of CD8^+^ T cell density and penetration into the tumor core. To further examine the CD8^+^ T cell regional distribution, tumor and stroma were spatially annotated on H&E sections as summarized in Fig. 1F. Using this approach, Fig. 1G shows a marked increase in CD8^+^ T cell distribution in both the tumor and stroma regions of NOS2^*Lo*^/COX2^*Lo*^ tumors when compared to NOS2^*Hi*^/COX2^*Hi*^ tumors. Fig. 1H-1I demonstrate sparse or stromal-restricted CD8^+^ T cells, respectively in NOS2^*Hi*^/COX2^*Hi*^ tumors. In contrast, Fig. 1J shows dramatically increased CD8^+^ T cell density and penetration into the tumor core in a NOS2^*Lo*^/COX2^*Lo*^ tumor. These results further implicate a regulatory role of elevated NOS2/COX2 tumor expression in abated CD8^+^ T cell penetration into the tumor core.

### Indomethacin Engages the Immune System for Long Term Tumor-Free Survival in Nos2^-/-^ Mice

Our earlier work has shown that murine 4T1 tumor cells can express high Nos2 and Cox2 levels and that Nos2/Cox2 blockade limited tumor growth (16, 17). Thus, 4T1 tumor bearing mice are representative of NOS2^*Hi*^/COX2^*Hi*^ breast tumors. Next, the influence of tumor Nos2/Cox2 expression on tumor immune status was explored in immunocompetent 4T1 tumor-bearing mice. We utilized indomethacin (INDO), a clinically available NSAID that has been shown to efficiently target tumors expressing COX2 due to the slow rate of release of INDO from the COX2 enzyme when compared to other NSAIDs such as aspirin (18, 19). In addition, INDO also increased expression of the PGE2 consumptive enzyme PGDH, unlike agents such as celecoxib (Supplemental Fig. 1). These key features of make INDO an ideal candidate for studying the impact of COX2/PGE2 on the disease progression of cancer. To avoid potential toxicity associated with the dual NOS inhibitor/NSAID combination, a Nos2^-/-^ mouse was utilized (20, 21).

When compared to untreated wild-type (WT) controls, reduced 4T1 tumor growth in Nos2^-/-^ mice, WT mice treated with INDO, or Nos2^-/-^ mice treated with INDO was previously observed (17). Herein, tumor-bearing Nos2^-/-^ mice exhibited a moderate reduction in tumor growth as described by substance enhancement ratio of 1.14 (SER: ratio of slopes of treated mice to untreated mice) for mice reaching tumor volumes of 500mm^3^, while INDO treatment was more effective at limiting tumor burden (SER 1.86). In contrast, Cox2 inhibition in the Nos2^-/-^ mouse markedly reduced tumor growth as defined by SER 5.59 where only 30% of mice were able to reach a tumor volume of 1000 mm^3^ (Supplemental Table 1). The remaining mice continued to show partial as well as complete responses as demonstrated by the survival curve shown in Fig. 2A. Further exploration revealed that Nos2/Cox2 blockade in the combination-treated mice could be categorized into groups of low and high responders to treatment (Fig. 2B-C), as well as cured mice (Fig. 2D). The animals with regressing tumors (Fig. 2D) were followed for an additional 15 days on Indomethacin and then a 30-day indomethacin-free period while in remission (Fig. 2A). These mice were rechallenged with 4T1 tumor cells in the ipsilateral mammary fat pad. The rechallenge tumors grew more slowly with 20% complete regression where the cured mice stayed in remission for five months or longer (Fig. 2A), which was consistent with our earlier work (17). These results suggest that resistance to tumor rechallenge was mediated by a robust immune response associated with Nos2/Cox2 blockade. Moreover, these results show a clear benefit of targeted Nos2/Cox2 inhibition leading to changes in the immune response and disease remission.

**Fig. 2.**
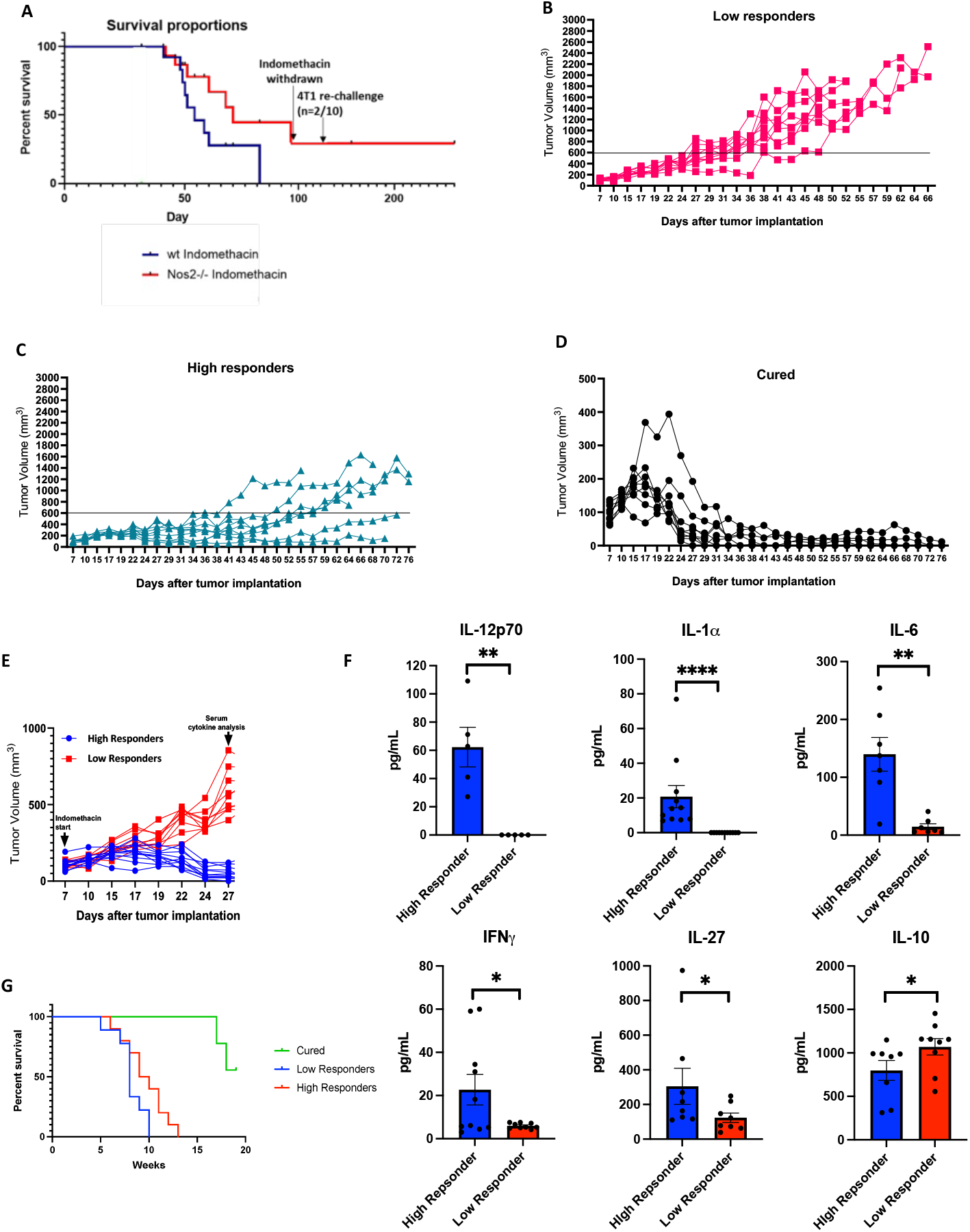
Orthotopic 4T1 tumor WT and NOS2^-/-^ mice treated with INDO. A) Survival analysis showing improved survival with INDO treatment; 20% of rechallenged mice were cured. Tumor growth curves for NOS2^-/-^ + INDO treated mice; B) mice with low treatment response, C) mice with high treatment response, and D) mice with regressing tumors. E) tumor growth in low vs high responders. F) Altered plasma cytokines in low vs high responders. G) survival curve.

To explore altered immune polarization in low and high responders to Nos2/Cox2 blockade, serum cytokine analysis was performed at day 27, reflecting an approximate time point when separation in tumor volume between low responding and high responding mice began (Fig. 2B,C,E). Serum cytokine analysis revealed considerably higher levels of proinflammatory Th1 cytokines including IL-12, IL-1α, IL-6, and IFNγ implicating a more robust systemic immune response in high responders to the Nos2/Cox2 blockade (Fig. 2F). Moreover, responding mice demonstrated increased IL-27, which suppresses FOXP3^+^ T_reg_ populations, thus providing additional supportive evidence that the high responders exhibit a more proinflammatory, anti-tumor immune microenvironment by limiting immunosuppressive Th2 immune polarization (Fig. 2F). In contrast, low responding mice produced low levels of IL-27 and proinflammatory cytokines, as well as higher levels of circulating immune suppressive Th2 cytokine IL10 (Fig. 2F). Importantly, survival analysis revealed improved survival in high responders and cured mice when compared to low responders (Fig. 2G).

### Nos2/Cox2 Blockade Alters Immune Polarization Within the TME

To further explore how Nos2/Cox2 inhibition alters the tumor immune microenvironment, flow cytometry (FACS), RNAseq gene expression, and multiplex imaging (CODEX) analyses were performed on 4T1 tumors harvested at eight days of INDO treatment. FACS and gene expression analyses revealed significant increases in CD45^+^ immune cells and CD11b^+^ myeloid cells in the INDO-treated tumors (Supplemental Fig. 2). FACs and CODEX imaging analysis revealed elevated CD3^+^, CD4^+^, CD8^+^ and CD19^+^ lymphocytes in WT+INDO and NOS2^-/-^+INDO treated mice, which supported increased T cell expression observed in NOS2^*Lo*^/COX2^*Lo*^ TNBC tumors (Fig. 3A-C, Supplemental Fig. 3, Fig. 1). Moreover, reduced CD25/CD4 ratios observed in all treated tumors suggest lower T_*reg*_ levels (Fig. 3D). CODEX analysis also showed increases in CD11c^+^CD169^+^CD8^+^ and CD11b^+^MHCII^+^ myeloid cells (Fig.3C) implicating a proinflammatory immune landscape. Additional supportive evidence of an antitumor immune landscape is provided by differential gene expression showing increased Th1 gene expression in INDO-treated mice, while higher CD19^+^ and CD20^+^ B cell gene expression was observed in Nos2^-/-^ mice (Fig.3D). Increased IFNγ, Gzmb, and Tbx21 as well as reduced IL10 expression were observed in INDO treated tumors, implicating cytolytic CD8^+^ T cell function (Fig. 3E). B cell gene expression profiles showed increased CD79 B-cell receptor and Ig-associated gene expression, while the B_regs_ marker CD1d1 was generally reduced, suggesting activated phenotypes (Fig. 3D). This is supported by CODEX analysis that showed increased CD19^+^CD38^+^ implicating mature, antibody producing B cell phenotypes (Fig. 3C). NOS2/COX2 blockade also promoted increased proinflammatory N1 neutrophil biomarkers including myeloperoxidase (*Mpo*), neutrophil elastase (*Elane*) and lactoferrin (*Ltf*) expression (Fig. 3D). Together these results show that NOS2/COX2 blockade promotes a proinflammatory, anti-tumor immune microenvironment involving increased N1/M1/Th1 antitumor immune phenotypes.

**Fig. 3.**
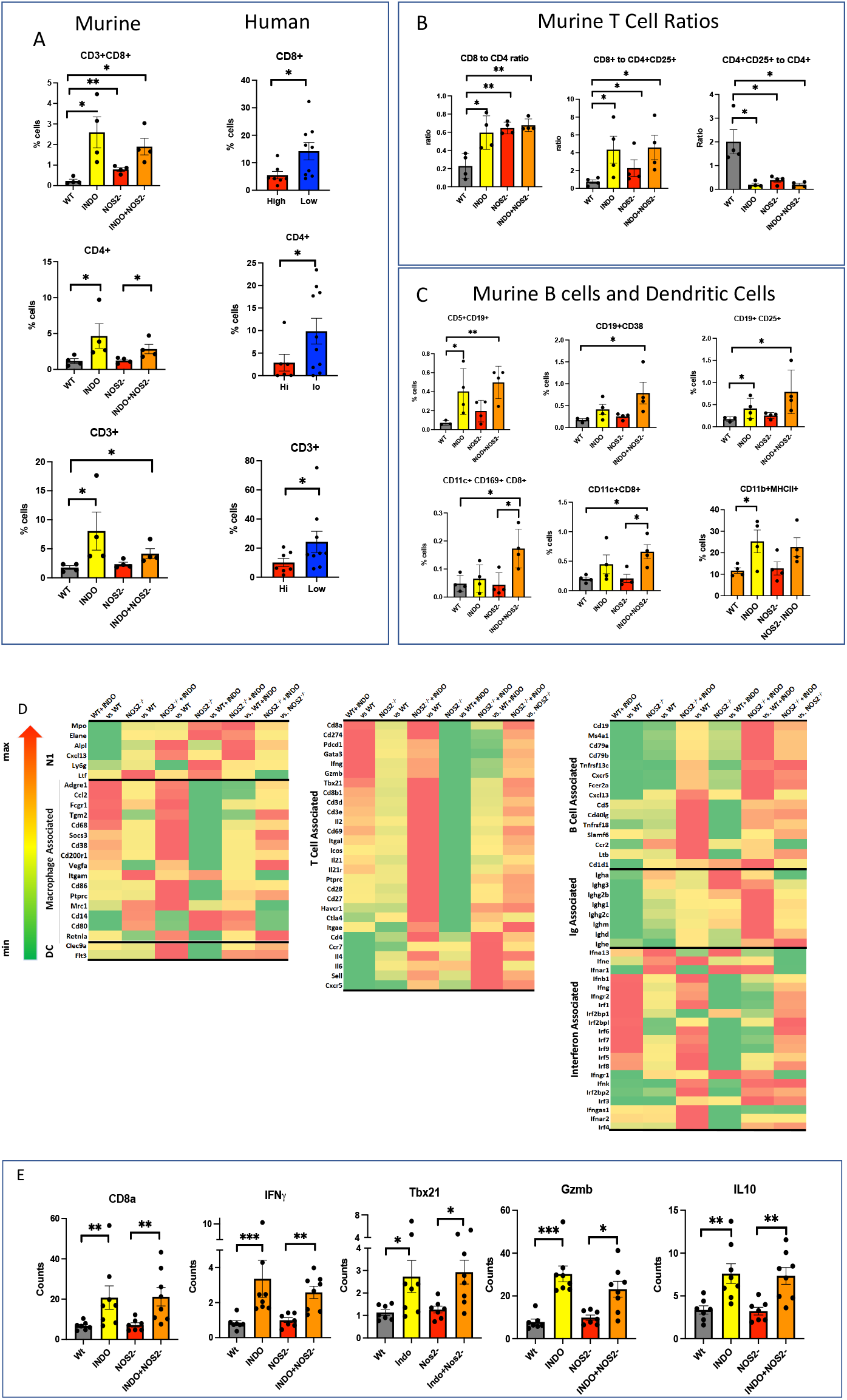
Flow cytometry, gene expression, and multiplex image analyses showing altered lymphoid populations. CODEX (mice) and Ultivue (patient TNBC tumors) analyses show A) increased T cells in both INDO-treated mice and NOS2^*Lo*^/COX2^*Lo*^ tumors. B) increased ratios comparing CD8^+^ to CD4^+^, CD8^+^ to CD4^+^CD25^+^, and CD4^+^to CD25^+^ T cell ratios. C) Increased B cells, dendritic cells, and proinflammatory macrophages. D) Gene expression heatmaps showing altered innate and adaptive biomarker expression. E) Gene expression analyses showing increased CD8a, IFNγ, TBX21, Gzmb, and IL10, which support increased cytolytic CD8^+^ T cells. (*p< 0.05, **p<0.01)

### Identification of Spatially Distinct Immune Phenotypes

Next, CODEX images were analyzed to assess the effects of Nos2/Cox2 blockade on the spatial distributions of immune cells in the 4T1 TNBC model. When compared to the untreated WT control, CODEX imaging revealed both increased density and penetration of CD8^+^ T cell populations into the tumor core in the INDO-treated tumors (Fig. 4A-D). Moreover, when compared to the untreated WT and Nos2^-/-^ tumors, increased CD8^+^ T cell clustering was observed in the INDO treated WT and Nos2^-/-^ tumors, which is supported by the spatial heatmaps shown in Fig. 4F,H (arrows), suggesting that cytolytic CD8^+^ T cells were markedly increased by Nos2/Cox2 blockade. Elevated B-cells (CD19^+^) as well as increased antigen presenting CD11c^+^ DC were also observed in the treated tumors (Fig. 4I-L). Taken together these results show that Nos2/Cox2 blockade promotes a robust immune response in 4T1 tumors, which is consistent with the immune microenvironment observed in NOS2^*Lo*^/COX2^*Lo*^ TNBC tumors (Fig. 1).

**Fig. 4.**
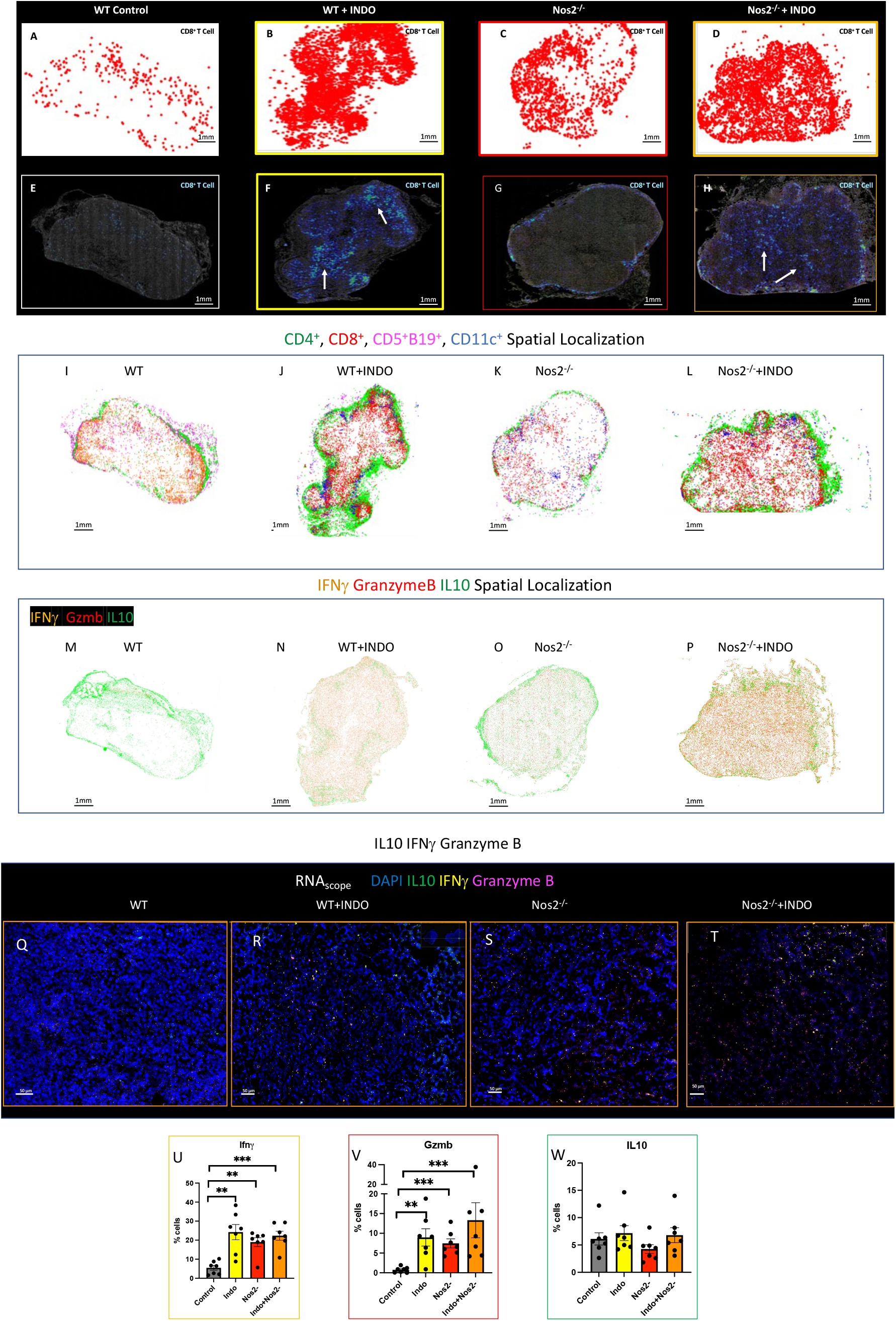
Nos2/Cox2 blockade alters lymphoid spatial localization and gene expression phenotypes. CD8^+^ T cell spatial orientation in A) untreated WT control, B) WT+ INDO, C) Nos2^-/-^, and D) Nos2^-/-^+ INDO where INDO treatments markedly increased CD8+ T cell density and penetration into the tumor core in INDO-treated mice. E-H) Heatmap analyses demonstrate enhanced CD8^+^ T cell clustering in INDO-treated mice. I-L) Distinct spatial orientation and layering of CD4^+^, CD8^+^, CD5^+^CD19^+^, CD11c^+^ lymphoid populations in control and treated tumors. M-T) RNA_*scope*_ analyses demonstrate altered IFNγ Gzmb, and IL10 expression in control and INDO-treated tumors. U-W) Quantification showing significantly elevated IFNγ and Gzmb in INDO-treated tumors, with no change in IL10 expession.

The density and spatial location of CD8^+^ T cells significantly improves clinical outcome as previously reported and shown in Fig. 1 and Fig. 4 (15). Increased tumor CD8^+^ T cell densities that penetrated deeply into the tumor epithelium predicted improved survival when compared to tumors resembling an immune desert or those with CD8^+^ T cell restriction to the tumor margin or stroma (15). Herein, untreated WT 4T1 tumors displayed immune deserts (Fig. 4) like those found in the NOS2^*Hi*^/COX2^*Hi*^ TNBC tumors (Fig. 1). To further characterize lymphoid populations in the 4T1 model, several immune phenotypes were examined. Spatial quantification of CODEX images demonstrated increased CD4^+^, CD8^+^, CD5^+^CD19^+^ and CD11c^+^ cells associated with Nos2/Cox2 blockade (Fig. 4I-L). In addition to CD11c^+^ APC cells, increased CD11c^+^CD169^+^CD8^+^ cells shown in Fig. 3C represent phenotypes that promote CD169-dependent antigen presentation, T cell priming and expansion that leads to CD8^+^ T cell polarization and cytolytic function (22, 23). Enhanced cytolytic T cells secrete IFNγ and granzyme b to facilitate tumor killing. Next, T cell polarization was examined using RNAscope, which confirmed enhanced *Ifnγ* and *Grzmb*, while *IL10* did not change in Nos2^-/-^ and INDO-treated mice (Fig. 4M-W). Mapping the expression in the whole tumor revealed a remarkable feature where *IL10* expression localized primarily on the tumor margins in all samples (Fig. 4M-P), residing with CD4^+^ T cells and B cells (Fig. 4I-L). In contrast, increased *Ifnγ* and *Gzmb* was spatially oriented in the tumor core (Fig.4Q-T) along with the CD8^+^ T cells (Fig. 4A-H). The spatial orientation of CD8^+^ T cells (Fig. 4A-H), *Ifnγ, Gzmb*, and *IL10* (Fig. 4M-W) provides additional evidence that Nos2/Cox2 inhibition leads to enhanced cytolytic T cell function occurring in the tumor core where it was surround by immunosuppressive conditions, which may be more conducive for the development of adaptive and humoral immunity. Together, these results suggest that Nos2/Cox2 blockade promotes increased APC populations and increased density and spatial distribution of CD8^+^ cytolytic T cells, that favor a fully inflamed anti-tumor immune microenvironment predictive of improved survival. Moreover, these observations suggest that the 4T1 tumor model provides an excellent platform for the evaluation of the impact of current and novel agents on immune polarization and therapeutic efficacy.

### Lymphoid Spatial Distribution Shows Unique Configurations

Further spatial examination of lymphoid structures demonstrated a layering effect of CD4^+^ and CD5^+^CD19^+^ cells, where the marginal area in WT tumors consists of densely populated CD4^+^ T cells and fewer CD5^+^CD19^+^ B cells surrounding CD11c^+^ dendritic cells and sparsely populated CD8^+^ cytolytic T cells. Treated tumors exhibit encapsulating CD4^+^ T cells with increased CD5^+^CD19^+^ B cells along the tumor margin and densely populated CD8^+^ cytolytic T cells that have infiltrated deeply into the tumor core (Fig. 4A-H). This configuration demonstrates distinct CD4^+^, CD8^+^, and B cell compartments, which are reminiscent of tertiary lymphoid structures (24). Gene expression analysis implicates elevated B cell receptor signaling and B cell activation markers with insignificant changes in B_*reg*_ markers that suggests proinflammatory phenotypes (Fig. 3). Moreover, increased CD19, CD20, IgH and IgM gene expression profiles suggest activated B cell phenotypes (Fig. 3E). Furthermore, CD19^+^, CD4^+^CD25^+^ and CD8^+^CD25^+^ cells were on the margins and not in the tumor core in WT+INDO and Nos2^-/-^+INDO treated tumors suggesting that the CD8^+^ cytolytic T cell response is spatially distinct (Fig. 3) This spatial orthogonality suggests that antitumor activity of the cytolytic CD8^+^ T cells is contained within the tumor core without effecting other regions of the tissue. In addition, B cells and CD4^+^ T cells activated in the TLS do not need to invade the tumor to elicit killing, which may at least in part account for the spatial localization of these cells in marginal regions that provides a spatial opportunity to maximize antitumor cross-talk. The pathway analysis summarized in Supplemental Fig. 4 shows strong recruitment of lymphoid cells, where the proximity of CD4^+^ T cells to CD19^+^ B cells may provide an important interaction during the development of adaptive immune response at the tumor edge. This could suggest that the margins have the potential for cross training of T cells and B cells. Together, these results provide further evidence that NOS2/COX2 blockade elicits a global proinflammatory immune spatial configuration that improves tumor eradication.

### Effect of NOS2/COX2 Inhibition on Taxol Therapeutic Efficacy

Given that NOS2/COX2 inhibition promotes a robust and spatially distinct antitumor immune response, its impact on therapeutic efficacy was examined. Docetaxel is a commonly used chemotherapeutic that often results in the development of drug-resistance (25). An earlier report demonstrated that neoadjuvant administration of a pan-NOS inhibitor (L-NMMA) with docetaxel improved therapeutic response in patients who were previously non-responsive to the taxane treatment (26). To better understand the effects of dual inhibition in combination with docetaxel, an *in vivo* study was performed in the 4T1 model (Fig. 5A). While docetaxel administered in two doses provided little effect on primary 4T1 tumor growth in WT mice, treatment with INDO significantly improved the response (Fig. 5B). Similarly, the same dosage administered to 4T1 tumor-bearing Nos2^-/-^ mice demonstrated significantly improved response by INDO treatment (Fig. 5B). Importantly, docetaxel+INDO treated Nos2^-/-^ mice exhibited significantly reduced lung metastatic burden (Fig. 5C) as well as reductions in severe side effects and co-morbidities associated with docetaxel+INDO treatment in WT mice. These results demonstrate the importance of Nos2/Cox2 blockade for improved response to docetaxel, which is consistent with a recently reported clinical study (14).

**Fig. 5.**
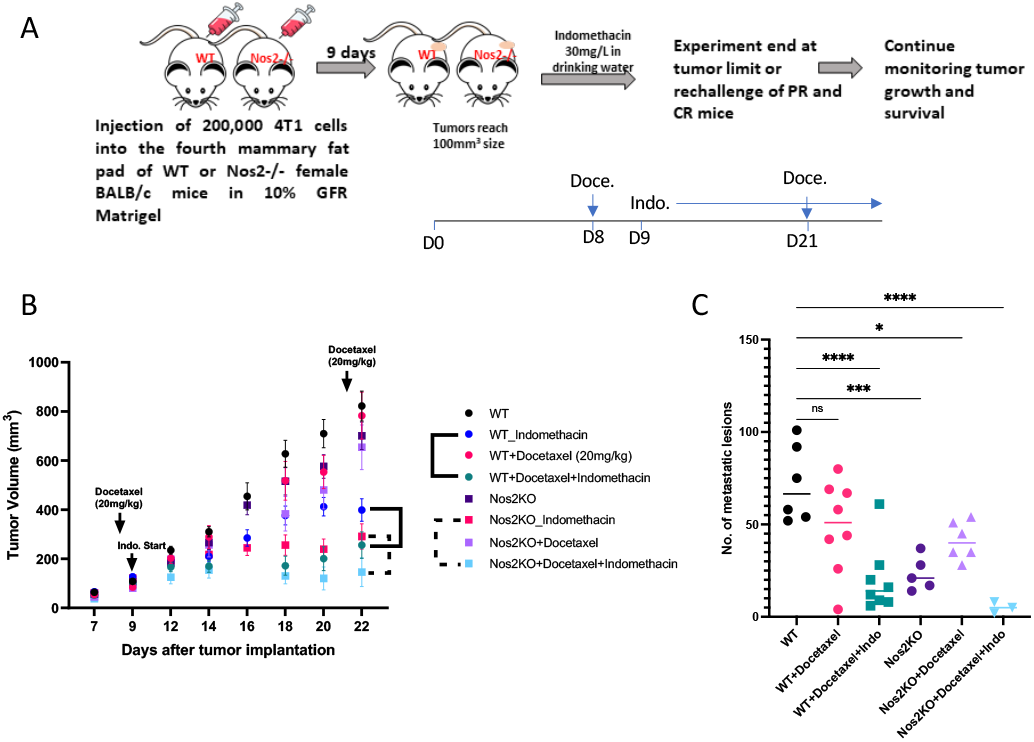
Nos2/Cox2 blockade improves docetaxel therapeutic efficacy. Treatment with **docetaxel** is summarized in panel A. B) Nos2/Cox2 inhibition reduces tumor growth and C) lung metastatic burden in taxane treated mice.

Metastasis is a key determinant of clinical outcome in cancer. Our earlier work showed that indomethacin played an important role in controlling growth of the primary tumor but had only a modest effect on lung metastatic burden. In contrast, dual Nos2/Cox2 blockade was able to significantly reduce metastatic lesions in 4T1 tumor-bearing mice (17). To explore the role of Nos2/Cox2 in metastatic development, INDO administration was initiated when 4T1 tumors reached a size of 100mm^3^ and then tumor resection/survival surgery was performed the following week. The mice were maintained for a total of 45 days and then euthanized, and lungs were harvested for evaluation of metastatic burden (Fig. 6 experimental design). Untreated WT mice exhibited large numbers of metastatic lesions while WT+INDO and Nos2^-/-^ mice had significantly reduced metastatic burden (Fig. 6C). Importantly, only one detectable metastatic lesion was observed among seven Nos2^-/-^+INDO-treated mice, which was less than 1% of the lesions observed in untreated WT mice (Fig. 6C,D). These results suggest that neoadjuvant Nos2/Cox2 blockade may provide a beneficial option for TNBC patients undergoing surgical resection.

**Fig. 6.**
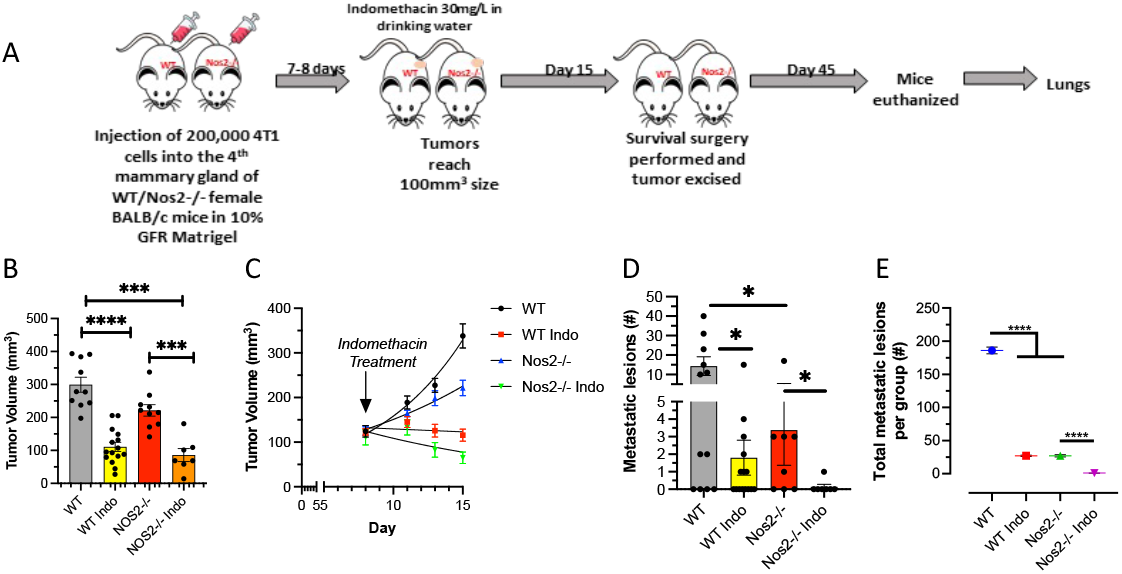
Nos2/Cox2 blockade improves surgery therapeutic efficacy. Survival surgery is summarized in panel A. B-C) Nos2/Cox2 inhibition reduces tumor growth and D-E) lung metastatic burden in treated mice.

## Discussion

The coinhibitory immune ligand programmed cell death ligand 1 (PD-L1; CD274) is elevated in TNBC. However, when compared to other cancers, these advanced tumors are less responsive to checkpoint inhibitors that target these ligands (27). In addition, advanced TNBC patients present with increased drug resistance due in part to the presence of immune suppressive cell types, making this disease more difficult to treat (28). Indeed, poor clinical outcome is, in part, due to elevated immunosuppressive tumor immune microenvironments. Herein, we show that elevated tumor expression of NOS2 and COX2 creates an immunosuppressive tumor immune microenvironment, as blockade of these pathways promotes proinflammatory immune polarization. Tumor NOS2 and/or COX2 overexpression occurs in numerous aggressive tumors including, lung, glioma, liver, pancreatic, cervical, gastric, esophageal, and ER-breast cancer. In ER-breast cancer patients, elevated NOS2 and COX2 tumor expression is a strong predictor of poor survival (4, 5, 7). Interestingly, p53 negatively regulates NOS2 and COX2 expression. Mutations in p53 are often associated with poor clinical outcome of aggressive tumors, which may in part lead to the elevated and chronic expression of NOS2 and COX2 (29, 30). Moreover, tumor NOS2 expression correlates with p53 mutation and predicts poor survival in ER-breast tumors (4). Herein, we show that elevated NOS2/COX2 signaling directly impacts both the immune response and spatial orientation of CD8^+^ T cells in TNBC. Importantly, NOS2/COX2 blockade markedly improved the immune profile and tumor infiltration of cytolytic CD8^+^ T cells, resulting in reduced tumor burden in 4T1 tumor-bearing mice. These results suggest that NOS2/COX2 overexpression promotes an immunosuppressive tumor immune microenvironment.

Within the TME, NOS2-derived NO and COX2-derived PGE2 can impact different aspects of the local immune response by tuning the area to favor an immunosuppressive microenvironment (10, 31, 32). Pro-oncogenic NO levels (100-300 nM flux) influence T cell function through the induction of apoptosis of CD4^+^/CD8^+^ Th1 phenotypes while favoring CD4^+^/CD8^+^ T_regs_ that produce the immunosuppressive cytokine IL-10 (12). In addition, these levels of NO activate TGFβ as well as IL-10 which prevents M1 macrophage development (33). Also, NO can inhibit immune infiltration of neutrophils, monocytes, and lymphoid cells through regulation of vascular adhesion molecules, such as ICAM, VCAM and MCP-1 thereby creating leukocyte deserts (34). Herein, the development of the immune architecture strongly shows several particularly strong changes in the immune profile in response to Nos2/Cox2 blockade including increased T cells, B cells, N1 neutrophils and dendritic cells (Fig. 3E). Where Cox2 inhibition by INDO in WT clearly effects the T cells populations and suppressed N1 neutrophil and B cells. In contrast, Nos2^-/-^ increases N1 and B cell infiltration (Fig. 3-4) suggests that Nos2/Cox2 blockade offers complimentary immune regulatory components. Analysis show INDO treated samples had higher levels *Ifng, Ifngr, Irf1, and Irf7* regardless of NOS2 status (Fig. 4). According to canonical pathway analysis summarized in Supplemental Fig. 4, Cox2 inhibition reprogrammed the downstream signaling cascade through IFNγ and activation of several T cell-related pathways including T Cell Receptor Signaling, Natural Killer Cell Signaling, ICOS-ICOSL Signaling in T Helper Cells, and Crosstalk between Dendritic Cells and Natural Killer Cells. In contrast, NOS2^-/-^ mice had increased B cell and N1 neutrophils. The dual expression T cell with B cell and N1 neutrophils in the combination appears to be important for the successful antitumor response.

When expressed by the tumor, PGE2 has a multidimensional impact on the overall tone of the immune microenvironment by favoring immunosuppression and anti-inflammatory mechanisms (10). Herein, we show that COX inhibition by indomethacin promotes proinflammatory/antitumor Th1 phenotypes of CD4^+^ and CD8^+^ T cells, which reduced tumor growth and metastatic burden and improved survival of 4T1 tumor-bearing mice. Like NO, PGE2 increases IL10 and TGFβ Th2 cytokines, which block DC maturation and antigen presentation (9, 10). In general, PGE2 suppresses Th1, cytolytic T cells and NK cells through cAMP-dependent mechanisms while attracting T_*reg*_ and MDSC immunosuppressive phenotypes (9, 10). Indeed, a key mechanism of immune suppression by PGE2 is its cAMP-dependent effect on T cells (10). Early studies have shown that PGE2 promotes T cell anergy and limited T cell responsiveness (35). Importantly, feedforward tumor NOS2/COX2 regulation augments NO/PGE2 levels in a synergistic manner leading to a chronically immunosuppressed tumor immune microenvironment (7).

Mature neutrophils play a key role in tumor eradication through increased MPO and reactive oxygen species (ROS) generation. Herein, we show that 4T1 tumors grown on a NOS2^-/-^ background in the presence of INDO increased the proportion of mature neutrophils in the tumor. In contrast, the higher proportion of immature neutrophils in the WT control, due to excessive GMC-SF production by the tumor promotes immunosuppression as well as setting the metastatic niche in the lung (36). While NO will drive many of the oncogenic pathways for EMT and migration, reduced pulmonary Ly6G was identified, which is important in seeding the metastatic niches in the lung (37). Moreover, primary tumors from NOS2^-/-^+INDO-treated mice exhibited elevated expression of neutrophil activation markers implicating limited immunosuppression and the promotion of antitumor neutrophil phenotypes by the dual treatment. This suggests a role for elevated ROS, which is known to be involved in tumor eradication (38). In contrast, NOS2-derived NO scavenges ROS directly, thus serving as an antioxidant, which protects the tumor (39). Importantly, elevated neutrophil infiltration identified in patients who responded to NOS inhibition/taxane therapy suggests that increased proportions of mature neutrophils is a key anti-tumor mechanism associated with NOS2 blockade in both mice and humans (14).

Another important feature of NOS2^-/-^+INDO combination therapy is increased B cell infiltration. The RNA_seq_ analysis identified the modulation of several B cell-associated genes. The RNAseq observations were further supported by FACS and CODEX analyses, which demonstrated an increase in B cell and APC density as well as reduced tumor growth and metastasis indicating a potentiated anti-tumor response. Importantly, these results are consistent with a recent clinical study, where chemo-resistant metaplastic breast cancer patients who responded to taxane and adjuvant NOS inhibition exhibited markedly increased B cell penetration into the tumor (14). Taken together, NOS2/COX2 blockade polarizes the tumor immune microenvironment favoring a potent proinflammatory, antitumor immune response.

The spatial localization of proinflammatory immune phenotypes is also influenced by NOS2/COX2 blockade, which led to dramatically increased cytolytic CD8^+^ T cell penetration deep into the tumor core. A layered effect was observed, reminiscent of an “Inside-out lymph node” consisting of cytolytic CD8^+^ T cells surrounded by separate layers of CD4^+^ T cells and B cells. Recently, tertiary lymphoid structures (TLS) have been detected in tumors, which correlate with improved survival (40, 41). This configuration suggests an orthogonality in the lymphoid population that is like lymph node structures. Furthermore, the spatial orientation of CD25^+^ T cells are on the margins and not in the cytolytic core, suggesting that NOS2/COX2 inhibition not only changes the immune tone but also the spatial arrangement of immune mediators. Importantly, this novel spatial orientation provides an opportunity for the T cells, CD11c^+^ DC, and B cells to cross-train each other leading to a more positive therapeutic response. A study describing an *in situ* vaccine utilizing an IL12 agent combined with radiation demonstrated that T cell-based antigen recognition along with humoral response through cross education was important for a stable resistance (42). In addition, elevated neutrophil density was observed in the necrotic cores of treated tumors. The spatial localization of antitumor neutrophils and cytolytic CD8^+^ T cells in the tumor core suggests a cooperative mechanism of these immune mediators (43). Given that neutrophils home to and lead CD8^+^ T cells into necrotic areas suggests a layering sequence of immune processing and activation (44). Together, the observed lymphoid layering may provide a unique and important spatial orientation associated with NOS2/COX2 targeted therapies that maximizes antitumor crosstalk between immune cells.

In summary, the therapeutic importance of a proinflammatory tumor environment indicates a need for therapeutic interventions that modulate tumor inflammation. The current study implicates tumor NOS2/COX2 expression as key immunosuppressive effectors that limit the efficacy of immune therapies. Importantly, the clinical availability of NOS2/COX2 targeting agents may provide a novel opportunity for therapeutically unresponsive tumors (14).

## Materials and Methods

### *In vivo* studies

Animal care was provided at the NCI-Frederick Animal Facility according to procedures outlined in the Guide for Care and Use of Laboratory Animals. Our facility is accredited by the Association for Accreditation of Laboratory Animal Care International and follows the Public Health Service Policy for the Care and Use of Laboratory Animals. Female BALB/c mice obtained from the Frederick Cancer Research and Development Center Animal Production Area were used for the *in vivo* studies and housed five per cage. Eight to ten-week-old female WT and Nos2^-/-^ BALB/c mice were shaved a day prior to tumor injection and then were injected subcutaneously into the fourth mammary fat pad with 200,000 4T1 TNBC cells. Tumor measurements began one week after tumor cell injection, using a Vernier caliper and calculated in cubic millimeter volumes according to the following equation

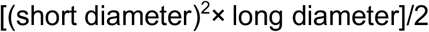

Upon reaching tumor size of 100mm^3^, tumor-bearing mice were divided into groups and treatment with 30 mg/L indomethacin in drinking water was initiated. The water was changed every MWF and treatment continued for the duration of the experiment unless otherwise specified. Tumors were harvested and snap-frozen and kept at -80°C until further use. Serial 10μm sections were cut for CODEX, RNAseq, and RNAscope analyses. For the combination treatment with docetaxel, taxane was administered at 20mg/kg of the drug in gel-chow (Nutra gel-chow, Bioserve) on day 8 and day 21 after tumor injection. For this study, Indomethacin treatment began on the following day and continued as mentioned previously. Experimental end point varied according to the study. Mice were euthanized at or before the tumors reach 2000mm^3^ size. For the survival surgery, tumors were resected at 500mm^3^ size. Upon ending the experiment, lungs were fixed in Bouin’s solution and metastatic lesions were counted. Survival studies associated with the effects of dual inhibition on immune response to tumor were also performed; indomethacin treatment was stopped in the mice achieving complete remission i.e., cured mice, and these mice were then injected subcutaneously in the fourth mammary fat pad on the opposite side with 200,000 4T1 cells. Tumor growth was monitored in these mice for a period of 5 months or until the tumor in the rechallenge site reached 2000mm^3^ size, whichever was earlier.

### Immunophenotyping of Tumors by Flow Cytometry

Mice were euthanized and tumors were collected and dissociated by mechanical dissociation (Miltenyi GentleMACS) in lysis buffer containing Collagenase and Dnase in 5% RPMI. Red blood cells were removed by incubating in ACK lysis buffer and washing with phosphate buffered saline (PBS). Cells were counted and equal numbers of cells were stained with the Live/Dead Aqua reagent (Amcyan) (1:1000) in PBS for 30 minutes followed by PBS wash and 20min at 4°C with Fc blocker (1:200) in Sorter buffer (1%FBS, 1mM EDTA in PBS). The cells were then stained with a panel of fluorophore-tagged antibodies against various immune cell markers including CD45-FITC, CD3-BV785, CD4-PECy7, CD8a-PerCPCy5.5, CD19-BV605, Tim3-APC, CD62L-PE, CD45-BV605, CD11b-PerCPCy5.5, CD11c-APCCy7, F4/80-APC, Ly6G-BV711, Ly6C-PECy7, CD206-FITC, PDL1-PE, MHCII (MHC IA/IE)-BV421. Samples were incubated for 20min at 4°C, washed and read on a flow cytometer. Respective unstained cells and FMO (Fluorescence Minus One) controls were used to set the positive gates during acquisition. Samples were acquired using the low/medium flow rate setting on the BD LSRII Sorp flow cytometer, normalized to tumor weight and analysis was performed using FlowJo software.

### Serum Cytokine Analysis

Blood was collected from mice by retroorbital bleed and saved in Microtainer^®^ Blood tubes (BD Biosciences, Cat. No. 365967). The vials were left undisturbed for 30 minutes at room temperature, spun down at 2000rpm for 10 minutes at RT and serum was collected, aliquoted and stored at -80°C. Samples were completely thawed and centrifuged prior to use. Multiple freeze-thaws were strictly avoided. Serum cytokine analysis was performed using LEGENDPlex Mouse Inflammation Panel from BioLegend (Cat. No. 740446) according to manufacturer’s instructions. This method employs the Sandwich ELISA principle. Briefly, 2x diluted serum samples were incubated on a shaker for 2h at RT with mixed beads tagged with antibodies against 13 different cytokines. The beads are distinguishable by size and the level of APC fluorophore on their surfaces. The beads were then washed and incubated with biotinylated-detection antibodies on a shaker for 1h at RT followed by a 30min incubation with Streptavidin-PE. Samples and standards were washed and read on the BD LSR Fortessa flow cytometer in the PE and APC channels and data was analyzed using the LEGENDPlex software.

### RNA Sequencing of Bulk Tumor

In brief, tissue samples were harvested after eight days of treatment with indomethacin and stored at -80°C. Two 10 μm slices were homogenized in the presence of TRIzol (ThermoFisher) and further purified with affinity column (RNeasy Mini Kit, Qiagen) following the manufacturer’s protocols. Extracted RNAs underwent RNA quality check in a bioanalyzer and only samples with a RIN (RNA Integrity Number) larger than 6 were used to make the RNAseq library prep. Sample libraries were prepped with the Illumina Stranded Total RNA Prep and paired-end sequencing performed according to the manufacturer protocol and sequenced in a NovaSeq 600 sequencing system. Reads of the samples were trimmed for adapters and low-quality bases using Cutadapt. Sequencing data were exported and then uploaded to the Partek Flow server for subsequent sample normalization and QC steps using the build-in RNAseq Data Analysis workflow. Differential expressed gene lists were generated with the Partek GSA algorithm which applies multiple statistical models to each individual gene in order to account for each gene’s varying responses to different experimental factors, and different data distributions. A 2-fold cutoff and p-value < 0.05 filter was applied to finalize the gene lists.

### CODEX^®^ Analysis

The CODEX protocol was performed according to Akoya User Manual, revision B.0. Square (22 × 22 mm) glass coverslips (72204-10, Electron Microscopy Sciences) were pre-treated with L-Lysine (Sigma, St. Louis, MO) overnight at room temperature. Coverslips were rinsed in distilled water, dried, and stored at room temperature. Fresh frozen tissue blocks were sectioned (10μm) on treated coverslips and stored in a coverslip storage box (Qintay, LLC) at -80°C until further use. CODEX antibodies, reagents (including those for conjugation of additional antibodies), and instrumentation were purchased from Akoya Biosciences (Marlborough, MA). Antibodies labeled for CODEX included CD279, CD86, Ki67, E-cadherin, CD19, PIMO, CD31, CD49f, vimentin, F4-80, alphaSMA, CD44v6, Ly6C, NOS2, CD206, CD25, CD11c, CD274, CD44, CD24, MHCII, CD3, CD90, CD5, CD71, CD45, CD4, CD169, CD38, CD8a, Ly6G, CD11b. Tissue sections were stained with an antibody cocktail consisting of 0.5-1μl of each antibody per tissue. CODEX assays were performed according to the manufacturer’s recommendations. Fluorescent oligonucleotide plates were prepared in black 96-well plates for image acquisition. Each CODEX cycle contains four fluorescent channels (three for antibody visualization and one for nuclear stain). For each cycle, up to three fluorescent oligonucleotides (5 μL each) were added to a final volume of 250 μL of plate buffer (containing Hoechst nuclear stain). For blank (empty) cycles, 5 μL of plate buffer was substituted for fluorescent oligonucleotides. Plates were sealed and kept at 4°C until use. For imaging, the CODEX coverslip was mounted onto a custom-designed plate holder and securely tightened onto the stage of a Keyence BZ-X810 inverted fluorescence microscope. Cycles of hybridization, buffer exchange, image acquisition, and stripping were then performed using an Akoya CODEX instrument. Briefly, that instrument performs hybridization of the fluorescent oligonucleotides in a hybridization buffer, imaging of tissues in CODEX buffer, and stripping of fluorescent oligonucleotides in the stripping buffer. CODEX multicycle automated tumor imaging of was performed using a CFI Plan Apo 20x/0.75 objective (Nikon). The multipoint function of the BZ-X viewer software (BZ-X ver. 1.3.2, Keyence) was manually programmed to align with the center of each tumor and set to 10 Z stacks. Nuclear stain (DAPI, 1:600 final concentration) was imaged in each cycle at an optimized exposure time of roughly 10 ms. The respective channels were imaged in the automated run using optimized exposure times. Raw TIFF images produced during image acquisition were processed using the CODEX image processer. The processer concatenates Z-stack images, performs drift compensation based on alignment of nuclear stain across images, and removes the out-of-focus light using the Microvolution deconvolution algorithm (Microvolution). The processer also corrects for non-uniform illumination and subtracts the background and artefacts using blank imaging cycles without fluorescent oligonucleotides. The output of this image processing was tiled images corresponding to all fluorescence channels and imaging cycles that were then visualized and analyzed using HALO software (Version 3.3.2541.383, Indica Labs Inc.). Segmentation of cells was performed using the nuclear channel and the cell cytoplasm was defined as a fixed width ring around each nucleus. Nuclear segmentation settings were optimized by visual verification of segmentation performance on random subsets of cells aiming to minimize the number of over segmentation, under segmentation, detected artefacts and missed cells. Cell type Annotation and Differential Marker Analysis Cell populations were gated as follows. All nucleated cells were first identified by positive nuclear signals. Cell phenotypes were defined based upon biomarker expression as judged by expert visual inspection.

### Immunohistochemical Analysis of Patient Tumor Sections: Ultivue^®^

Formalin-fixed paraffin embedded (FFPE) tissue sectioned at 4 μm and mounted on SuperFrost Plus slides were stained with a FixVUE Immuno-8™ Kit (formerly referred to as UltiMapper® kits, Ultivue Inc., Cambridge, MA, USA; CD8, PD-1, PD-L1, CD68, CD3, CD8, FoxP3, and panCK/SOX10 cocktail) using the antibody conjugated DNA-barcoded multiplexed immunofluorescence (mIF) method (1). These kits include the required buffers and reagents to run the assays: antibody diluent, pre-amplification mix, amplification enzyme and buffer, fluorescent probes and corresponding buffer, and nuclear counterstain reagent. Hematoxylin and Eosin (H&E) and mIF staining was performed using the Leica Biosystems BOND RX autostainer. Before performing the mIF staining, FFPE tissue sections were baked vertically at 60–65 °C for 30 min to remove excess paraffin prior to loading on the BOND RX. The BOND RX was used to stain the slides with the recommended FixVUE (UltiMapper) protocol. During assay setup, the reagents from the kit were prepared and loaded onto the autostainer in Leica Titration containers. Solutions for epitope retrieval (ER2, Leica Biosystems cat# AR9640), BOND Wash (Leica Biosystems cat# AR9590), along with all other BOND RX bulk reagents were purchased from Leica). During this assay, the sample was first incubated with a mixture of all 8 antibody conjugates, next the DNA barcodes of each target were simultaneously amplified to improve the sensitivity of the assay. Fluorescent probes conjugated with complemental DNA barcodes were then added to the sample to bind and label the first round of 4 targets; a Round 1 fluorescent image was then acquired. Next, a gentle signal removal step was used to remove the fluorescent probes of the first set of markers before adding the fluorescent probes specific for the second set of 4 markers before imaging the slide a second time to acquire the Round 2 fluorescent image. There was no need for quenching, bleaching or other means to minimize signal between rounds. Before each round of imaging, the stained slides were mounted in Prolong Gold Anti-Fade mountant (Thermo Fisher Scientific, cat# P36965 and coverslipped (Fisherbrand Cover Glass 22 × 40mm, #1.5).Digital immunofluorescence images were scanned at 20× magnification. Round 1 and 2 images were co-registered and stacked with Ultivue’s UltiStacker software. The Immuno8 images used the following marker/fluorophore combinations: FITC (CD8 Round 1 (R1), CD3 Round 2(R2)), TRITC (PD-1 R1,CD4 R2), Cy5 (PD-L1 R1, FoxP3 R2), Cy7 (CD68 R1, panCK/Sox10 R2) and the custom kit used the following combinations: FITC (Alpha SMA R1, CD68/CD163 R2), TRITC (Arg-1 R1, CD15 R2), Cy5 (iNOS R1, IDO1 R2), Cy7 (CD20 R1, CD14 R2). The digital images were then analyzed using HALO™ software (45).

### Statistical analysis

One-way ANOVA with Tukey’s multiple comparisons test or Mann-Whitney test was employed to assess statistical significance using the GraphPad Prism software. Significance is reported as *p ≤ 0.05, **p ≤ 0.01, ***p ≤ 0.001, ****p≤0.0001.

## Fig. Legends

**Supplemental Table I.** Substance Enhancement Ratio (SER) Summary of control and INDO-treated tumors.

|  | Days from injection to<br>500mm<sup>3</sup> | % mice<br>reaching<br>500mm<sup>3</sup> | SER | Days from<br>injection to<br>1000mm<sup>3</sup> | % mice reaching<br>1000mm<sup>3</sup> | SER |
| --- | --- | --- | --- | --- | --- | --- |
| WT | 13-18 | 80% | 1 | 23-30 | 100% | 1 |
| Nos2<sup>-/-</sup> | 13-18 | 70% | 1.143 | 23-30 | 100% | 1 |
| WT INDO | 27-38 | 90% | 1.864 | 34-46 | 90% | 1.7 |
| Nos2<sup>-/-</sup> INDO | 27-38 | 30% | 5.59 | 34-46 | 30% | 5.1 |

**Supplemental Fig. 1.**
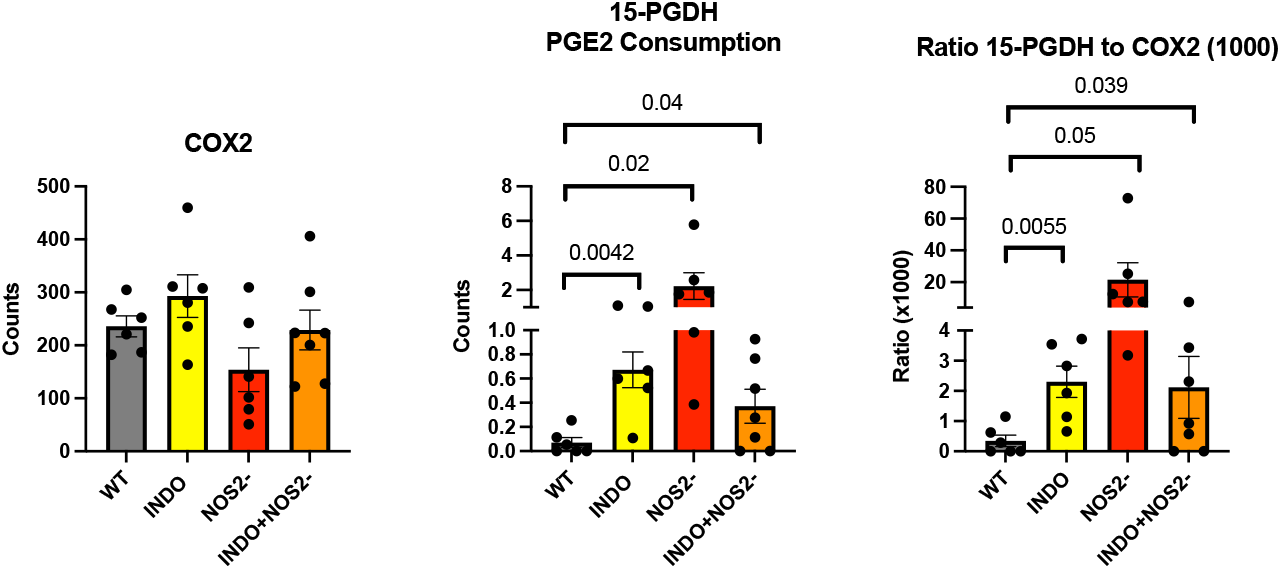
**Expression of** A) Cox2, B) the PGE2 consuming enzyme 15-PGDH C) the15-PGDH/Cox2 ratio. (p-values are shown in Fig.).

**Supplemental Fig. 2.**
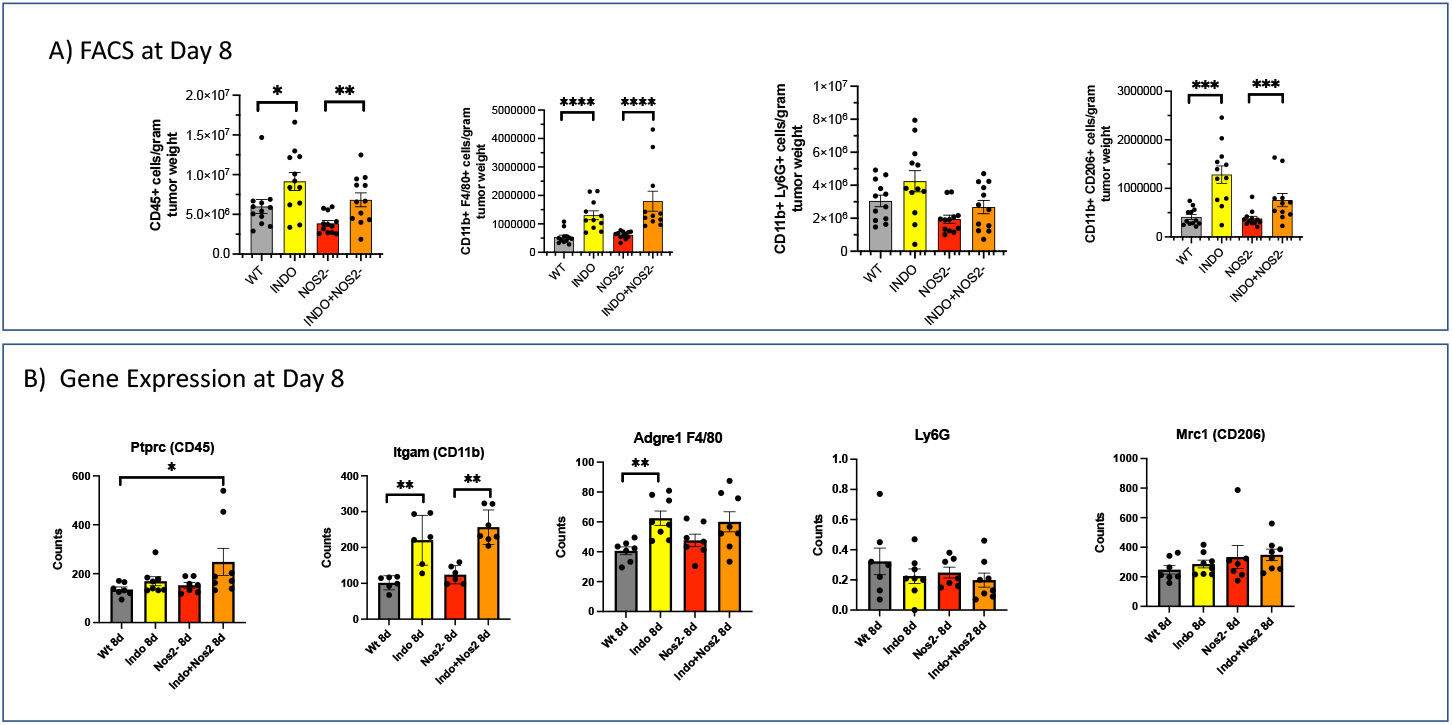
FACs analysis and RNA seq gene expression show similar patterns of characteristic myeloid markers. *p< 0.5, **p<0.01

**Supplemental Fig. 3.**
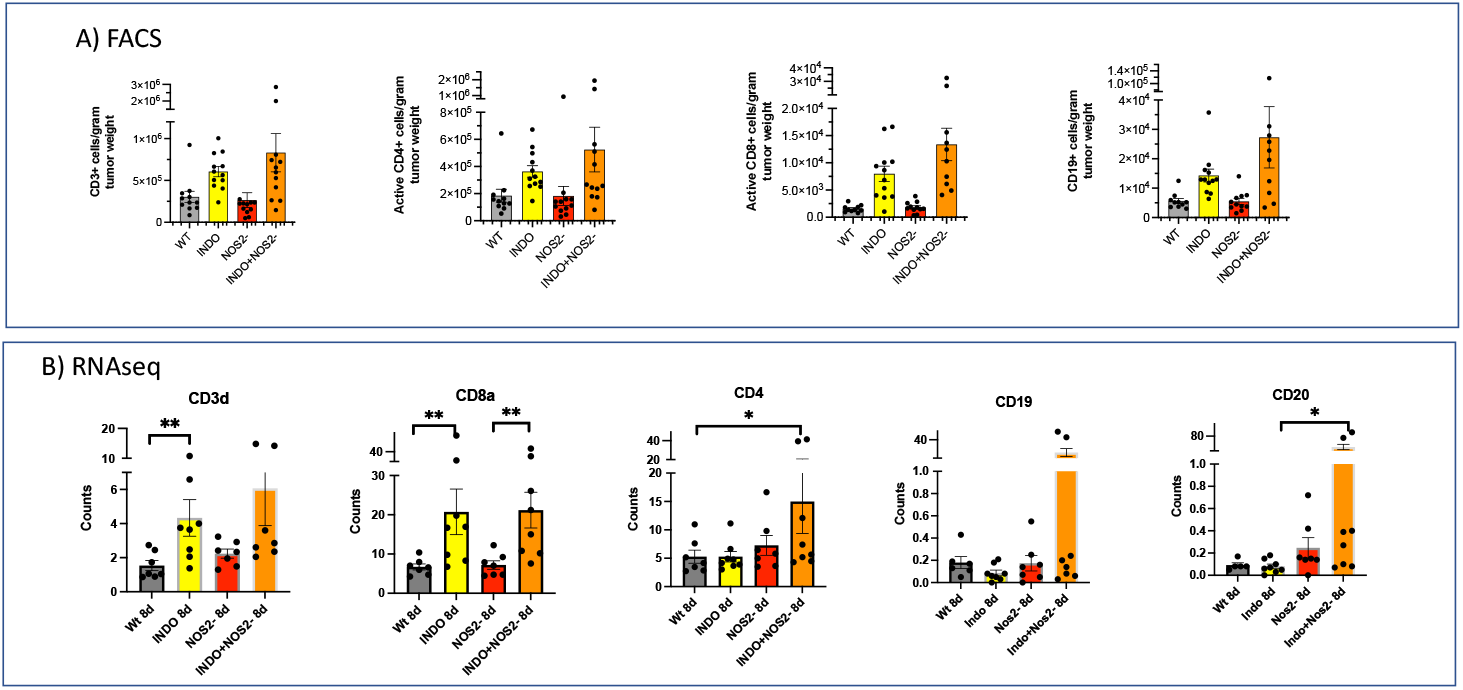
FACs analysis and RNA seq gene expression show similar patterns of lymphoid markers. *p< 0.5, **p<0.01.

**Supplemental Fig. 4:**
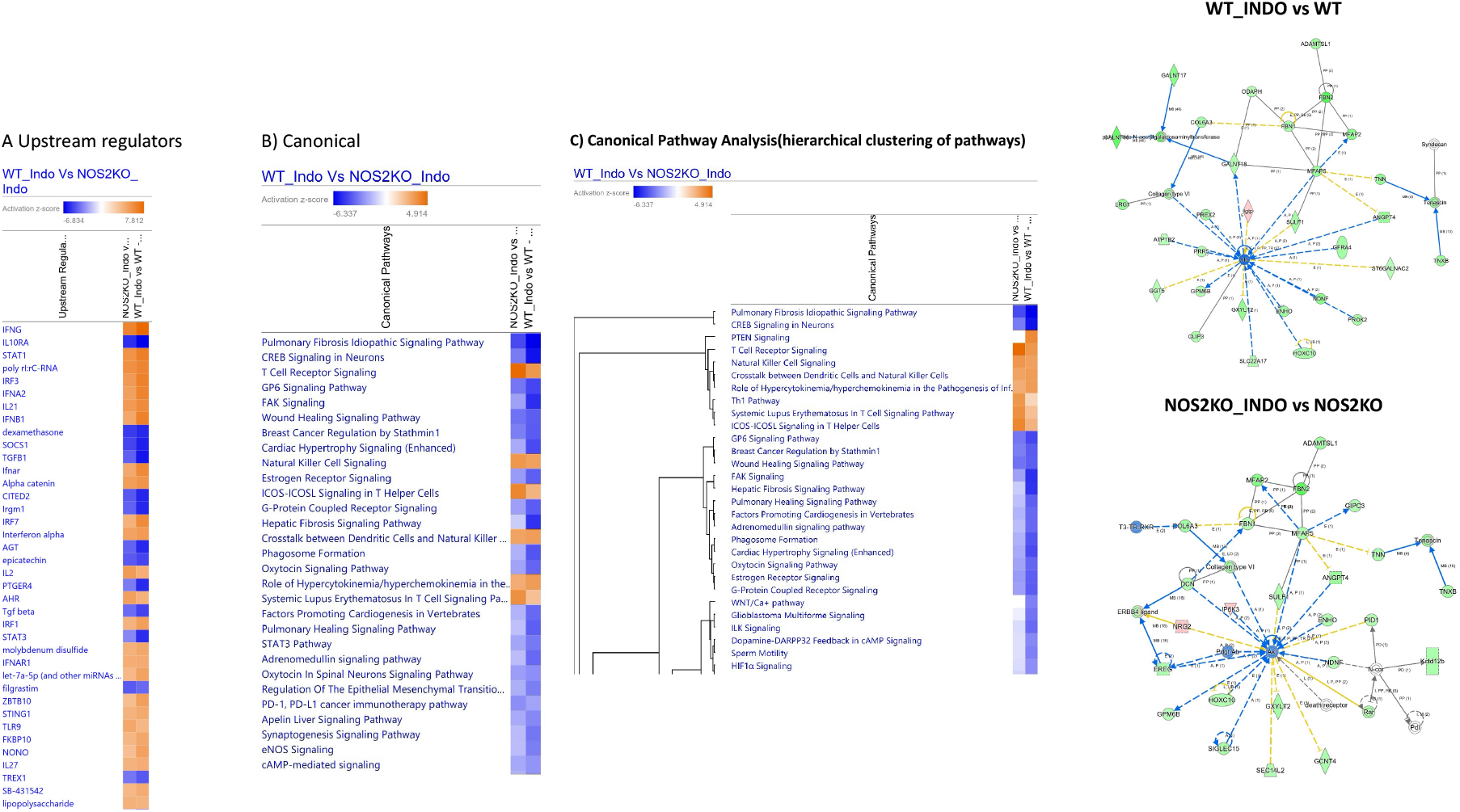
Pathway Analysis.

